# SUSTAINED DOPAMINERGIC PLATEAUS AND NORADRENERGIC DEPRESSIONS MEDIATE DISSOCIABLE ASPECTS OF EXPLOITATIVE STATES

**DOI:** 10.1101/822650

**Authors:** Aaron C. Koralek, Rui M. Costa

## Abstract

We are constantly faced with the trade-off between exploiting actions with known outcomes and exploring alternative actions whose outcomes may be better. This balance has been hypothesized to rely on dopaminergic neurons of the substantia nigra pars compacta (SNc)^1^ and noradrenergic neurons of the locus coeruleus (LC)^2–3^. We developed a behavioral paradigm to capture exploitative and exploratory states, and imaged calcium dynamics in genetically-identified dopaminergic SNc neurons and noradrenergic LC neurons during state transitions. During exploitative states, characterized by motivated repetition of the same action choice, we found dichotomous changes in baseline activity in SNc and LC, with SNc showing higher and LC showing lower sustained activity. These sustained neural states emerged from the accumulation of lengthened positive responses and hysteretic dynamics in SNc networks, and lengthened negative responses in LC. Sustained activity could not be explained by classical reinforcement learning parameters, and in SNc but not LC, emerged in subpopulations coding for response vigor. Manipulating the sustained activity of SNc and LC revealed that dopaminergic activity primarily mediates engagement and motivation, whereas noradrenergic activity modulates action selection. These data uncover the emergence of sustained neural states in dopaminergic and noradrenergic networks that mediate dissociable aspects of exploitative bouts.

---

At any given moment, animals must choose their next action from a vast repertoire of possible behavioral responses. Some actions have been performed repeatedly in the past and therefore have well-known outcomes, while others have less certain but potentially better outcomes. In addition, there are fluctuations in the motivational drive to perform some actions over others, depending on the current state of both the environment and the animal. This trade-off between exploiting known actions (low choice entropy) and exploring alternative ones (high choice entropy) has been proposed to rely on midbrain dopaminergic neurons of the substantia nigra pars compacta (SNc)^1^ and noradrenergic neurons of the locus coeruleus (LC)^2–3^. Deficits in choice reversal learning^4^ and attentional set switching^5^ have been demonstrated following dopamine (DA) depletion, and computational modeling work has predicted a central role for DA signaling in modifying action selection probabilities^6^. Similarly, recent work suggests that levels of norepinephrine (NE), specifically the noradrenergic projections to prefrontal cortices, modulate levels of stochastic responding in rodents^7–8^. Both dopaminergic and noradrenergic systems therefore appear to play a central role in motivating and structuring adaptive behavior.

Although past work has studied isolated exploitative and exploratory choices^9–13^, the majority of this work has focused on single-trial decisions, thus obscuring the longer-term state changes that define exploitative and exploratory states of action selection. These exploitative states, characterized by engaged and motivated performance of the same well-learned action to achieve a desired outcome, might be similar to what is colloquially referred to as “being in the zone” or the “hot hand effect”^14–15^. However, although both the motivational and repeated choice aspects of exploitative states go hand in hand, it is unclear if they are mediated by the same neural substrates.

We developed a novel behavioral paradigm in mice that probes action selection among many possible actions over long time scales and allows us to bias behavior towards exploitative or exploratory states using environmental changes. This paradigm permitted us to study the behavior of animals away from ceiling or floor performance, and hence to study the emergence of bouts of exploitative choices, i.e. periods of motivated performance of the same actions. We imaged the activity of populations of individual dopamine neurons of the SNc and noradrenergic neurons of the LC and found striking changes in sustained dopaminergic and noradrenergic activity that cumulatively emerged when animals were in exploitative behavioral states. These exploitative states were marked by lengthened response plateaus and hysteretic network dynamics in SNc neurons, as well as lengthened response depressions in LC neurons. The sustained activity of SNc, but not LC, neurons was related to the vigor or motivation of the behavior. Finally, we induced sustained changes in the excitability of SNc dopaminergic neurons and LC noradrenergic neurons and found that these systems subserve dissociable aspects of action motivation and selection, with SNc mediating the motivation to engage in action bouts and LC mediating choice entropy. These data reveal that these two major neuromodulatory systems display sustained neural states that mediate different aspects of exploitative behavioral states.

## Sustained Dopaminergic and Noradrenergic Modulations in a Novel Task for Probing Exploitative and Exploratory Behavioral States

To develop a framework for studying exploitative and exploratory states in mice, we created a nose poke sequence task in which mice can choose between many possible actions. Mice were placed in an operant chamber with 3 equidistant nose pokes (**Fig. 1a**). A sequence of 3 pokes in a specific order was rewarded. Importantly, mice were given no trial structure and few cues to guide learning, but instead had to actively explore the environment to determine the reward structure. When mice performed the target sequence, reward was supplied via a central reward port. There are 27 possible sequences, providing a broad distribution of potentially selectable actions, and the entropy of this distribution captures levels of exploitative and exploratory behavior

**Figure 1.**
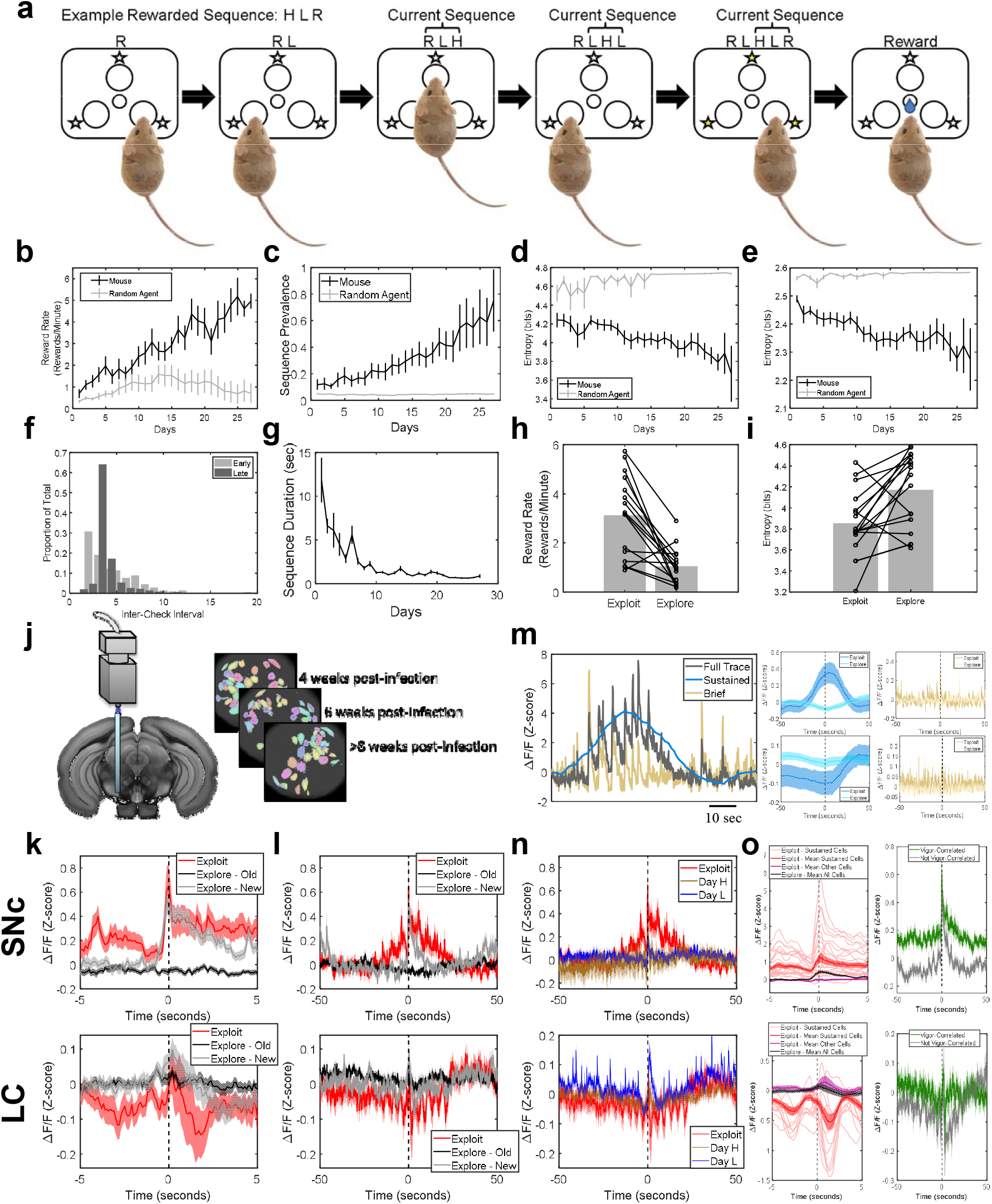
Sustained dopaminergic and noradrenergic activity modulations in a novel task for probing action motivation and selection. **a.** Task schematic. Mice are presented with three equidistant nose poke ports and must discover a rewarded sequence of three nose pokes in order. There are no trials, but instead a moving buffer of the last 3 nose pokes is monitored for the rewarded sequence. Over the course of training, **b.** reward rate increases, **c.** the prevalence of the rewarded sequence increases, **d.** the entropy of the distribution of selected actions decreases, and **e.** the entropy of the transitions between pokes decreases. Gray lines indicate estimates of chance performance. **f.** Early in learning, mice check for reward after most response pokes, but late in learning, they begin to perform 3 response pokes before checking for reward. **g.** The average time necessary to complete the rewarded sequence decreases with training. **h.** Reward rate decreases and **i.** entropy increases following a change in the rewarded sequence. **j.** Schematic of endoscope imaging. **k-l.** PSTHs time-locked to exploitative rewards before the sequence change (“Exploit”; red), perseverative errors after the sequence change (“Explore-Old”; black), and exploratory rewards after the sequence change (“Explore-New”; gray) in SNc (top row) and LC (bottom row) with a short (k) or long (l) time axis. **m.** Left: Schematic showing full trace (black), slowly-varying component (blue), and quickly-varying component (tan). Right: PSTHs time-locked to reward for SNc (top row) and LC (bottom row) using the slowly-varying components (left column, blue) or quickly-varying components (right column, tan). **n.** PSTHs of activity when all sequences are rewarded with high (80%; brown) or low (20%; blue) probability relative to effects seen in exploitative states (red) in SNc (top) or in LC (bottom). **o.** Left: PSTHs of individual neurons that exhibit changes in sustained levels of activity during exploitative states (pink) in SNc (top) and LC (bottom) time-locked to performance of the rewarded sequence. The mean activity of other neurons in the network during exploitative states (purple) and the mean activity of all cells during exploratory states (black) are shown for comparison. Right: Mean activity of vigor-related neurons (green) and non-vigor-related neurons (gray) time-locked to reward achievement during exploitative states. Error bars denote s.e.m.

Performance improved significantly over training, as seen both in an increase in reward rate (**Fig. 1b**) and an increase in the proportion of pokes that compose the rewarded sequence relative to total pokes (**Fig. 1c**). Chance levels of performance were assessed by modeling an agent that performs the same number of pokes as the mice on each day, but selects each poke randomly (**Fig. 1b-e**, gray lines). Animals performed significantly above chance level for all behavioral measures. Importantly, after training, mice were well above floor performance but also below ceiling performance, allowing us to study the transitions between exploitative and exploratory states. Over the course of training, we observed a decrease in the entropy of the animals’ selected sequences (**Fig. 1d**), as well as a decrease in the entropy of the animals’ selected transitions between nose pokes (**Fig. 1e**), suggesting that animals are initially sampling a relatively wide range of possible actions, but gradually refine these choices to focus more on the rewarded sequence. When examining the number of pokes at response ports between checks for reward at the reward port (“Inter-Check Interval”), we observed that animals check for reward after a majority of response pokes in early learning, but begin to structure behavior into groups of three nose pokes in late learning (**Fig. 1f**), suggesting that they have learned to mostly perform three-poke sequences. The time that mice took to perform the rewarded sequence also decreased significantly with training (**Fig. 1g**).

When the animals were proficient at performing a target sequence, we changed the target sequence to be rewarded. This sequence change occurred within a behavioral session (**Supp. Fig. 1**). Across all animals, we observed a significant decrease in performance (**Fig. 1h**) and an increase in entropy **(Fig. 1i**) immediately following the rule change, suggesting that changing the reward structure of the environment was successfully driving animals into a more exploratory state.

We next imaged calcium dynamics in genetically-identified dopaminergic and noradrenergic cells of the SNc and LC, respectively, through chronically-implanted gradient index (GRIN) lenses (**Fig. 1j**). We first examined phasic bursting in the populations before and after a change in the rewarded sequence (**Fig. 1k**), with a focus on three conditions. Namely, “Exploit” designates the epoch before the rule change when mice were exploiting a well-known reward structure, with peri-stimulus time histograms (PSTHs) time-locked to reward delivery following performance of the target sequence. The “Explore” conditions, in contrast, designate the epoch after the rule change when mice were exploring a novel reward structure, and this is subdivided into “Explore-Old”, representing perseverative errors when mice performed the previously-rewarded action that was no longer rewarded, and “Explore-New”, representing trials when mice performed the newly-rewarded action. Importantly, sequence changes occurred within a behavioral session, ensuring that levels of fluorophore expression and bleaching were comparable across all conditions. In both regions, we observed qualitatively similar phasic responses to rewards during exploitative and exploratory epochs. However, these phasic responses arose from different baseline levels of activity, with SNc baseline activity enhanced, and LC baseline activity reduced, during exploitation (**Fig. 1k**). In SNc, exploitative and exploratory rewards resulted in comparable peak magnitudes, despite the change in baseline activity, consistent with recent reports suggesting that reward expectation is marked by increased baseline activity rather than decreased peak amplitude^16^. Expanding the time axis, we found that these baseline changes developed slowly across multiple trials, lasting for roughly 60 seconds surrounding exploitative rewards (**Fig. 1l**).

We first asked whether these baseline changes were due to the averaging of many brief, staggered bursts of activity or were instead a slowly-varying component of the activity. To disambiguate these possibilities, we preprocessed our calcium data in a manner that separates quickly-varying and slowly-varying components of the signal (**Fig. 1m** and Methods) and we then used these component traces for our analyses. For both SNc and LC, we found that the changes in baseline activity were not observed when using only the quickly-varying components (**Fig. 1m**, right, tan), suggesting that the observed baseline shifts are not due to the averaging of many brief, jittered bursts. However, the sustained effects were still present when using only the slowly-varying components of the activity (**Fig. 1m**, right, blue).

We next asked whether these sustained activity changes were due simply to differences in reward rate during exploitation and exploration. We therefore ran animals on a version of the task in which all possible three-poke sequences were rewarded with either high probability (“Day H”; 80%) or low probability (“Day L”; 20%). We did not observe changes in baseline activity in either SNc or LC on either Day H or Day L (**Fig. 1n**). Importantly, the reward rate on Day H was comparable to that during exploitation, but the entropy was significantly higher (**Supp. Fig. 2**), suggesting that the baseline shifts were more closely related to changes in choice entropy than they were to changes in reward rate. In addition, we did not observe sustained changes in baseline activity in early learning (**Supp. Fig. 3**), or in dopaminergic neurons of the ventral tegmental area (VTA; **Supp. Fig. 4**). This phenomenon was also not the result of averaging across neurons. We found that roughly 20-25% of individual neurons in SNc and LC exhibit these sustained changes in baseline activity (**Supp. Fig. 5a**), with clearly distinct activity profiles relative to other neurons in the network during exploitative states or to all neurons during exploratory states (**Fig. 1o**).

Due to the slowly-varying nature of these baseline changes, we wondered whether they could be representing behavioral variables in a reinforcement learning (RL) context. We therefore fit a basic RL model to our behavioral data to extract estimates of action value (Q), state value (V), and reward prediction error (RPE). We found that the overall peri-event correlation of activity in all neurons and the different RL variables was strikingly low (**Supp. Fig. 6a-b**). In addition, we constructed reward-locked PSTHs using the estimates of Q, V, and RPE, and found these to exhibit very little activity at time points distant from reward (**Supp. Fig. 6c**). Finally, we correlated the full time-courses of Q, V, and RPE with smoothed fluorescence traces from SNc and found these correlations to also be very weak (**Supp. Fig. 6d**). We found, as expected, that individual cells vary greatly in the strength and direction of their correlations with these RL parameter estimates (**Supp. Fig. 6e**), but these correlations did not explain sustained reward-locked activity during exploitative relative to exploratory epochs (**Supp. Fig. 6f**). Conversely, we also found no strong relationship between the cells that we classified as exhibiting sustained effects (in **Fig. 1o**) and the distribution of their correlations with these RL parameter estimates (mean±SD, Q: 0.198±0.295, V: 0.142±0.299, RPE: 0.059±0.249). Together, these results suggest that, although the activity of many cells in the network was correlated with classical RL parameters, the observed changes in sustained activity cannot be accounted for simply by slowly-changing estimates of action and state value or RPE. Therefore, we examined if neurons exhibiting baseline shifts would correspond to neurons previously found in SNc related to movement initiation and vigor, which are distinct from those responding to reward^17^. We found that neuronal responses in both regions aligned more closely with movement initiation (**Supp. Fig. 5b**) than with reward achievement (**Fig. 1k**). When we separated neurons based on the correlation of their activity with movement vigor (see Methods), we found that vigor-correlated SNc neurons had significantly higher baseline activity during exploitation than other recorded neurons (**Fig. 1o**, right). Interestingly, the sustained activity decreases observed in LC neurons were not seen in vigor-correlated LC subpopulations, but instead in non-vigor-correlated LC subpopulations (**Fig. 1o**, right). These data suggest that the observed baseline shifts in SNc occur preferentially in neurons whose activity reflects the motivation to perform actions, but not in LC which has previously been implicated in action choice^2,7–8^.

## Sustained Activity Cumulatively Emerges in Dopaminergic and Noradrenergic Networks During Exploitative Action-Reward Bouts

Although we found that the sustained activity changes were not due to average reward rate (**Fig. 1n**), we wondered whether the reward structure changed following the sequence change. We therefore performed a “reward autocorrelation”, where time 0 indicates the occurrence of a reward and the occurrence of other rewards is averaged time-locked to this point. We found no change in the overall temporal structure of reward occurrence before or after the sequence change (**Fig. 2a**). Given that the SNc neurons that showed sustained activity seemed to be related to the motivation to perform an action, we investigated if there were periods of repetition of the actions that lead to reward (similar to the “hot hand effect”) during exploitative states. We therefore defined “target action-reward bouts” (hereafter “action-reward bouts”) as clusters of action-reward pairs (low choice entropy) that are separated from each other by less than 10 seconds and separated from other action-reward pairs by more than 20 seconds, and we then analyzed activity based on the action-reward pair’s position within a bout (**Fig. 2b** and **Supp. Fig. 7a**). Because the inter-event timing within action-reward bouts is not required to be constant, we first investigated whether the number of action-reward bouts changed across behavioral states, despite previously observing no shift in the overall temporal pattern of action-reward events (**Fig. 2a**). We found that the rate of occurrence of action-reward bouts increased significantly in exploitative relative to exploratory states (**Fig. 2c**), suggesting that action-reward bouts could be an important characteristic of the transitions between these behavioral states. In addition, animals responded significantly faster within action-reward bouts relative to other times (**Supp. Fig. 7**), suggesting that these bouts represent periods of enhanced vigor and motivation to perform the rewarded action. We therefore investigated neural responses throughout these bouts during exploitation and we found that dopaminergic activity accumulated over the course of a bout (**Fig. 2d**, top), while LC activity decreased consistently over the course of the bout (**Fig. 2d**, bottom). These patterns were not present during exploration (**Fig. 2e**), on Day H (**Fig. 2f**), or on Day L (**Fig. 2g**). Importantly, bouts of reward are common on Day H; however, they do not correspond to repetition of the same target action on Day H, but rather to performance of bouts of different actions. Therefore, the “hot hand effect”, or motivated repetition of the same action, should be minimal during this manipulation in natural settings, as many actions lead to reward.

**Figure 2.**
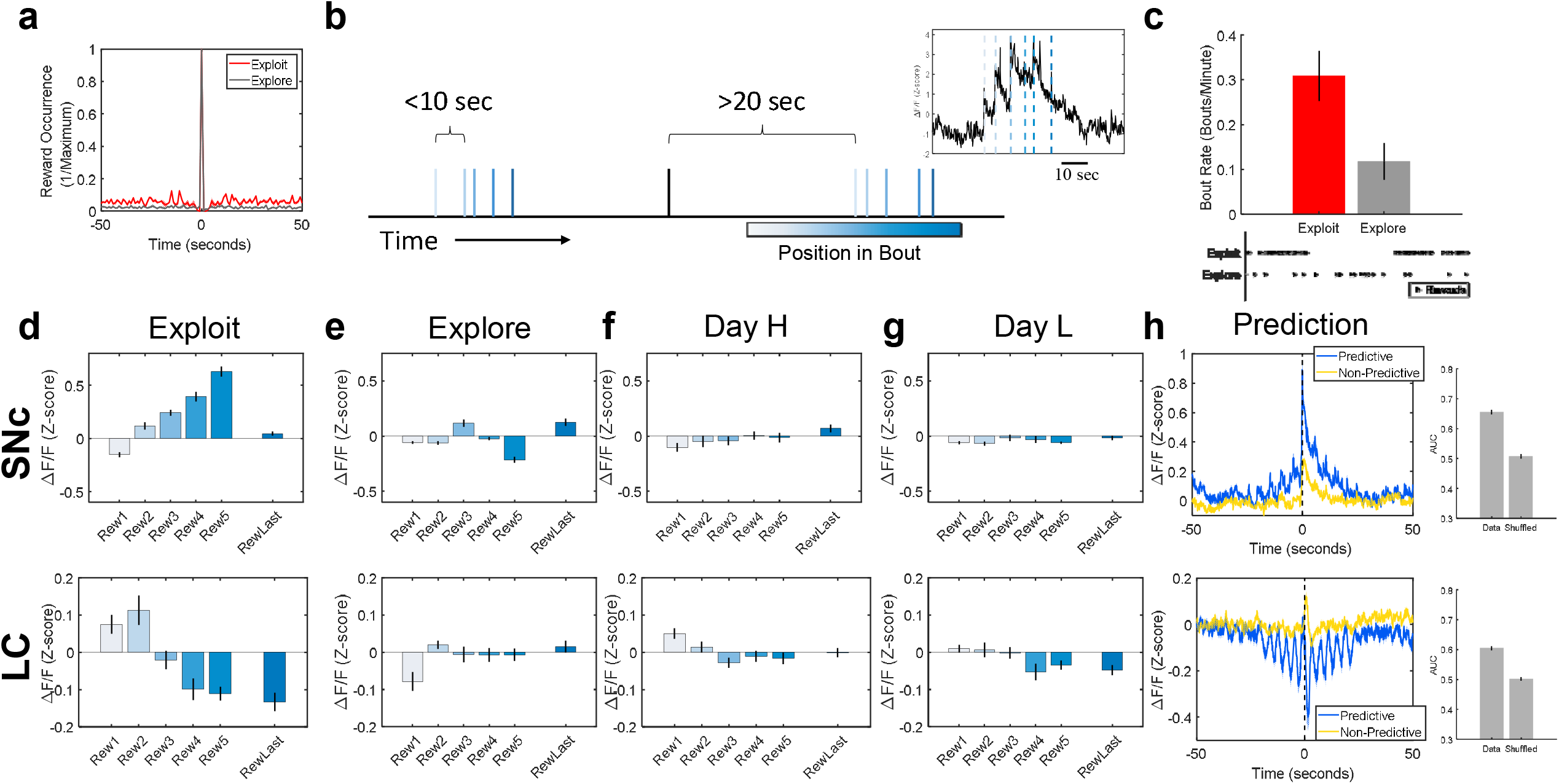
Activity accumulates positively in SNc and negatively in LC during exploitative action-reward bouts. **a.** Average occurrence of action-reward events time-locked to the occurrence of other action-reward events. There is no change in the mean temporal profile of action-reward event occurrences between exploitative (red) and exploratory (gray) states. **b.** Action-reward bouts are defined as clusters of action-reward pairs separated from each other by less than 10 seconds and separated from other action-reward pairs by more than 20 seconds. **c.** The raw number of action-reward bouts is increased in exploitative (red) relative to exploratory (gray) states. Bottom: Example raster showing occurrences of action-reward pairs clustering more into bouts during exploitation relative to exploration. **d-g.** Mean baseline activity preceding action-reward events separated by position within action-reward bouts in SNc (top) and LC (bottom) during exploitative states (c), during exploratory states (d), on Day H (e), and on Day L (f), **h.** Left: Neurons in SNc (top) and LC (bottom) whose activity is predictive (blue) and non-predictive (yellow) of reward bouts using a wiener filter. Right: Area-under-the-curve assessment of wiener filter performance using activity from individual SNc (top) or LC (bottom) cells relative to prediction performance in cases in which the category labels were shuffled. Error bars denote s.e.m.

We next asked whether the activity profile we observed during action-reward bouts was sufficiently characteristic to enable prediction of bout occurrences from neural activity. We found that prediction of action-reward bouts by single neurons was significantly better than chance (**Fig. 2h**, right). Furthermore, if we considered only neurons whose activity was most predictive of action-reward bouts, we found that these predictive neurons exhibited stronger sustained changes in activity surrounding exploitative action-reward pairs than the rest of the population in both SNc and LC (**Fig. 2h**). Together, these data suggest that sustained activity accumulates positively in SNc and negatively in LC as animals perform bouts of the same target action sequence in exploitative, but not exploratory, states. Furthermore, these bouts characterize an exploitative behavioral state, whereby animals are engaged in repeatedly performing a well-known action to achieve a favorable outcome.

## Altered Neuronal Response Dynamics Drive Sustained Activity

We next asked what neuronal response differences during exploitative and exploratory states could produce the distinct ways in which activity accumulates over the course of action-reward bouts to produce sustained activity shifts. We therefore quantified the average length of all positive and negative neural activity transients during exploitative and exploratory epochs. Remarkably, we found an increase in the duration of positive response transients in SNc neurons during exploitative relative to exploratory behavioral states, resulting in extended response plateaus (**Fig. 3a**). Importantly, this increase in duration of response plateaus was not explained by changes in response magnitude (**Fig. 3a,b**) and there was no change in the duration of negative response transients. In addition, the lengthened positive response transients in SNc neurons during exploitative states were not observed during early learning (**Supp. Fig. 3**), on Day H or Day L (**Supp. Fig. 2**), in the VTA (**Supp. Fig. 4**), or in the subset of SNc neurons whose activity was positively correlated with RL parameter estimates (**Supp. Fig. 8**). In contrast, in LC neurons, we found no change in positive response transients across these behavioral states, but the duration of negative response transients was significantly longer in exploitative relative to exploratory states, resulting in extended response depressions (**Fig. 3b**). Response transients from both regions were also fit individually with exponential functions, and with this metric we again found an increase in the duration of positive transients in SNc and negative transients in LC during exploitative states, with no associated increase in the magnitude of transients (**Fig. 3c,d**). To examine whether these activity changes in individual neurons were also reflected in changing network interactions, we analyzed the network correlation structure. Surprisingly, we found activity in SNc cells to be more correlated across the network during action-reward bouts specifically during exploitative states, with smaller correlations during exploratory states or outside of action-reward bouts in exploitative states (**Fig. 3e**). LC cells were more correlated with each other during action-reward bouts relative to non-bouts in both exploitative and exploratory states (**Fig. 3g**). To generate a more granular view of the consequences of these dynamics, we created cross-correlation histograms, where activity from all cells was time-locked to large fluorescence bursts in other simultaneously-recorded cells. Importantly, in the SNc during exploitative states, we found that cells in the network tended to increase activity together, but then continued to fire afterwards, exhibiting network-level hysteretic effects (**Fig. 3f**). In LC, correlated activity was generally increased during exploitation, but the shape of this response was unchanged (**Fig. 3h**). The asymmetry seen in SNc during exploitative states was also not present in early learning (**Supp. Fig. 3**), on Day H or Day L (**Supp. Fig. 2**), in the VTA (**Supp. Fig. 4**), or in the subset of SNc neurons whose activity was positively correlated with RL parameters (**Supp. Fig. 8**). Dopaminergic and noradrenergic networks therefore both exhibit striking changes in response dynamics across exploitative and exploratory behavioral states, with dopaminergic populations also entering a regime of hysteretic network interactions.

**Figure 3.**
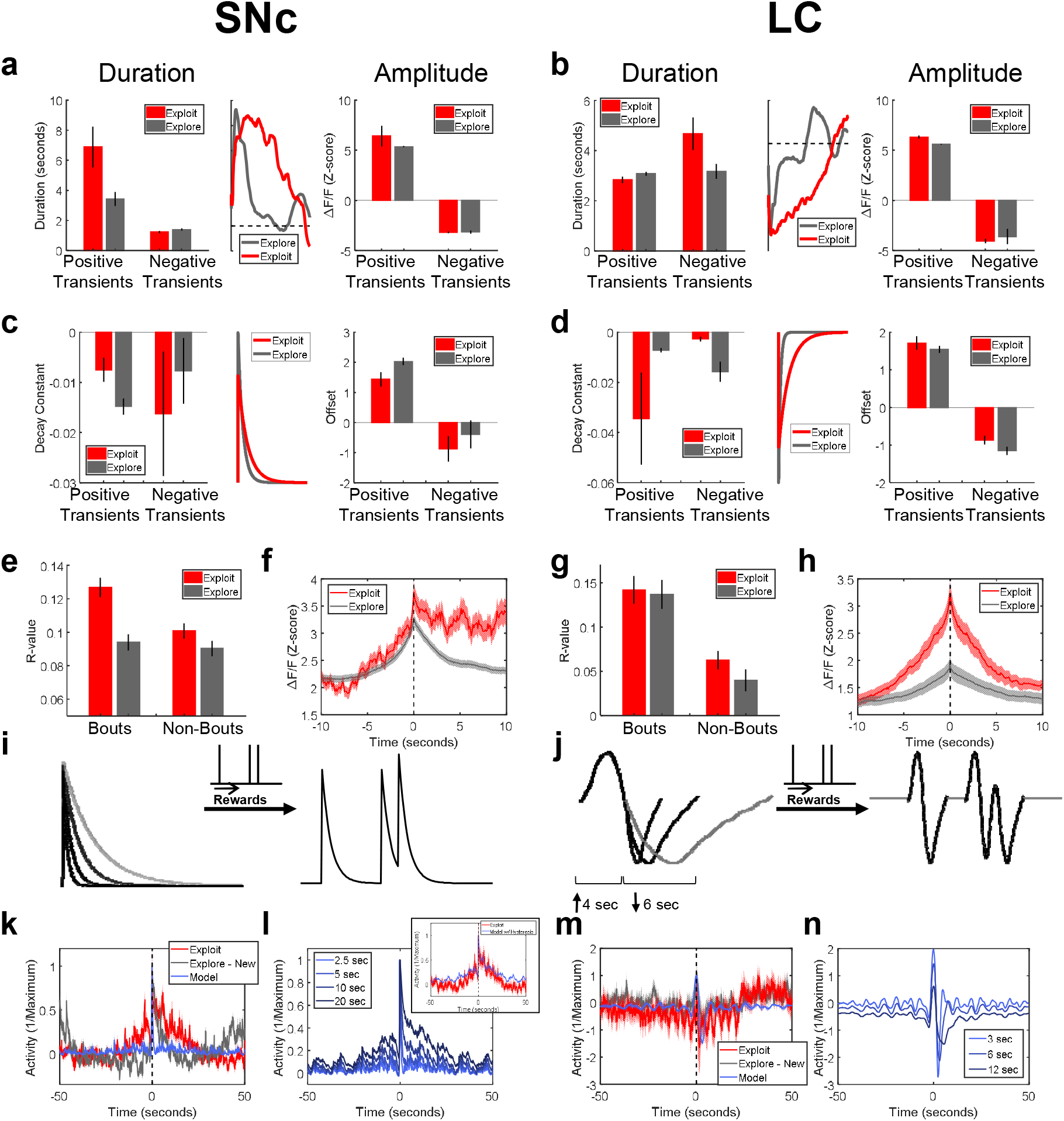
Extended response plateaus in SNc and depressions in LC produce the sustained accumulation of activity. **a-d.** Mean durations (left), amplitudes (right), and example traces (middle) of positive and negative transients in SNc (a,c), and in LC (b,d) calculated directly from response transients (a,b) or from exponential fits to response transients (c,d) during exploitative (red) or exploratory (gray) states. **e,g.** Mean peri-event activity correlations in or out of action-reward bouts during exploitative (red) or exploratory (gray) states in SNc (e) or LC (g). **f,h.** Cross-correlation histogram of neuronal activity time-locked to large fluorescence bursts in other cells during exploitative (red) and exploratory (gray) states in SNc (f) and LC (h). **i-j.** Schematics showing IRFs used to produce reward convolution traces for SNc (i) and LC (j). **k-n.** Reward convolution traces with typical IRFs in SNc (k) and LC (m), and with varying length IRFs in SNc (l) and LC (n). Model responses in blue, data for comparison in red and gray. Inset (l): Reward convolution traces with 10 second IRF and including hysteretic dynamics. Error bars denote s.e.m.

To investigate whether these changing response dynamics could produce the observed changes in sustained activity, we created convolution traces by convolving an impulse response function (IRF) with the occurrences of action-reward events in our behavioral data (**Fig. 3i,j**). For SNc, this IRF was a simple exponential of varying length (**Fig. 3i**), while for LC, this IRF was the smoothed average population response to unexpected rewards (**Fig. 3j**). As we increased the duration of the IRFs, we observed the emergence of sustained activity surrounding reward that matched that observed in SNc during exploitative states (**Fig. 3l,n**). This was not observed with IRF durations more closely matched to the statistics of our neural data in baseline settings (**Fig. 3k,m**). However, for IRFs matched to our data, the addition of hysteretic network dynamics to the model resulted in responses nearly identical to those seen in our data during exploitative states (**Fig. 3l**, inset; **Supp Fig. 9**). The LC IRF, on the other hand, is biphasic, with both a positive and negative phase that are asymmetric in baseline settings (**Fig. 3j**). If we alter the length of the negative phase of this IRF while holding the positive phase constant in our convolution model, we again see the emergence of sustained baseline changes with longer IRFs (**Fig. 3n**). Taken together, these data suggest that the increased duration of positive transients (response plateaus) and network hysteresis in SNc, together with the increased duration of negative transients (response depressions) in LC, can recapitulate the sustained activity observed during exploitative states.

## Increasing Dopaminergic or Noradrenergic Excitability Differentially Modulates the Motivation and Selection of Ongoing Actions

We next asked whether sustained changes in baseline activity levels could play a causal role in shifting between exploratory states and exploitative states, marked by action-reward bouts. Because these baseline shifts were due to changes in neural response dynamics that accumulate over the course of exploitative bouts, we used chemogenetic manipulations (hM3Dq) to enhance the excitability of genetically-identified dopaminergic and noradrenergic populations^19–20^ (**Fig. 4a**, rather than optogenetics to briefly drive excitability).

**Figure 4.**
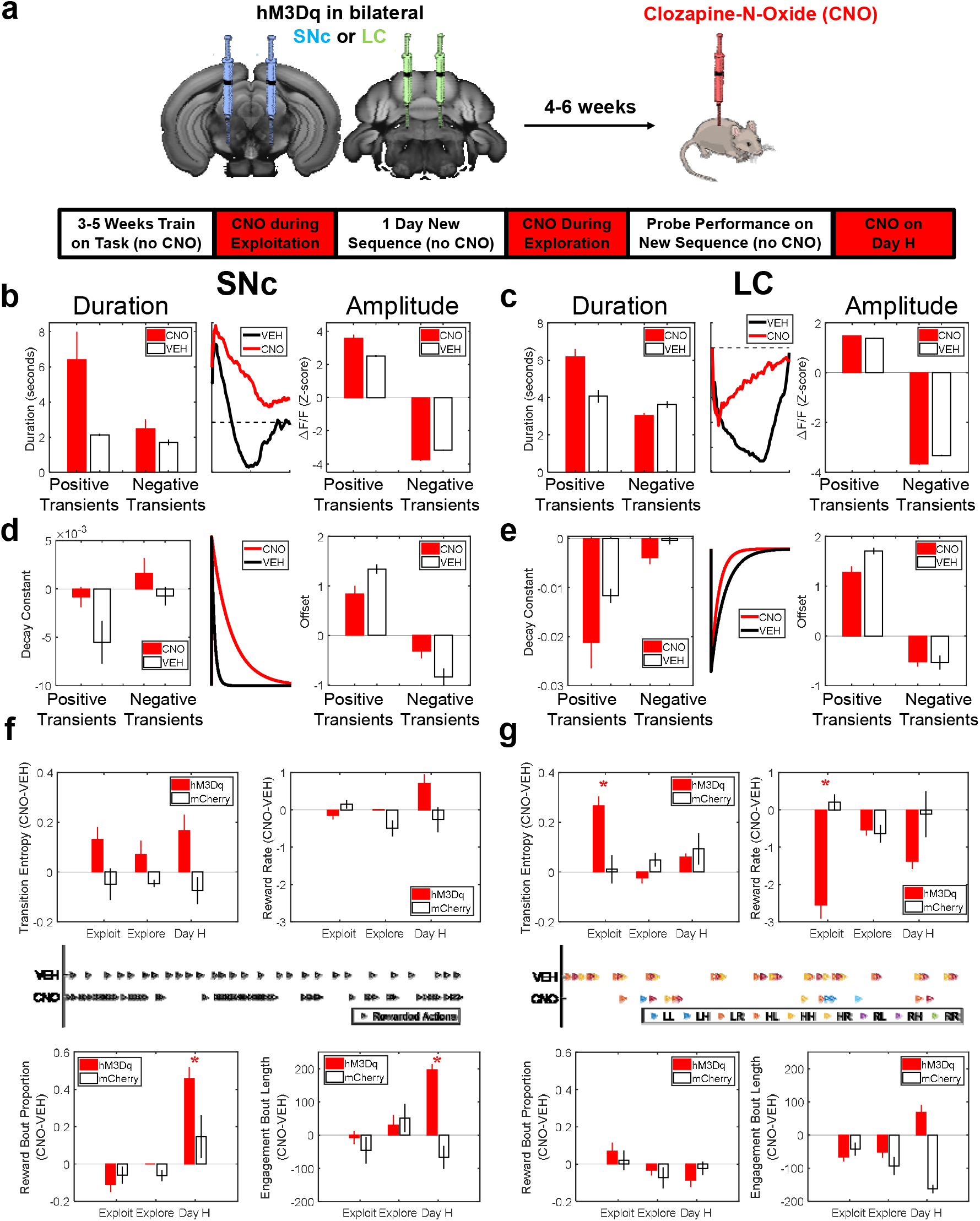
Modulating excitability in dopaminergic and noradrenergic neurons differentially biases action motivation and selection. **a.** Experimental timeline. **b-e.** Mean durations (left), amplitudes (right), and example traces (middle) of positive and negative transients in SNc (b,d), and in LC (c,e) calculated directly from response transients (b,c) or from exponential fits to response transients (d,e) following injection of CNO (red) or VEH (black). **f.** Transition entropy (top left), reward rate (top right), proportion of action-reward pairs that occured in bouts (bottom left), and duration of engagement bouts (bottom right) in animals expressing hM3Dq in SNc (red) and animals expressing mCherry in SNc (black), represented as the difference between values seen following CNO or following VEH (CNO-VEH). Center: Example raster showing the timing and clustering of action-reward events following injections of VEH (top) or CNO (bottom). **g.** Same measures as (f), but for animals expressing hM3Dq in LC (red) and animals expressing mCherry in LC (black). Center: Example raster showing the selected transitions between nose pokes following injections of VEH (top) or CNO (bottom). Error bars denote s.e.m. Asterisks denote significance at p<0.05.

Animals expressing either hM3Dq or mCherry were first imaged following intraperitoneal injections of either clozapine-N-oxide (CNO; the hM3Dq ligand) or vehicle (VEH) to determine the neuronal changes associated with chemogenetic activation. Following administration of CNO, we found a lengthening of the duration of positive response transients in both SNc (**Fig. 4b,d**) and LC (**Fig. 4c,e**) relative to VEH, as well as a shortening of the duration of negative response transients in LC.

We therefore trained animals on the task and noted the effects of CNO administration during exploitative states, exploratory states, and on Day H. Exploitative action-reward bouts involve both the motivation to perform actions, similar to the “hot hand effect”, as well as the selection of the same action to perform. Because sustained activity in SNc was related to response vigor and sustained activity in LC was not, we hypothesized that the motivation and selection aspects of exploitative states might be differentially mediated by dopaminergic and noradrenergic systems, respectively. Indeed, during exploitative states, we found that enhancing LC excitability with CNO produced an increase in transition entropy and a decrease in reward rate relative to VEH, and this effect was not seen in control animals expressing mCherry (**Fig. 4g**, top row, and **Supp. Fig. 10**). The same LC manipulation did not produce any effect during exploratory states, when animals still do not know the correct action sequence, showing that this was not a general disruption of behavior. Conversely, we found no change in transition entropy or reward rate when enhancing SNc excitability (**Fig. 4f**, top row, and **Supp. Fig. 10**), suggesting that the selection of an appropriate range of actions to perform is mediated by the noradrenergic, but not dopaminergic, system. We therefore asked whether enhancing SNc excitability instead affected the motivation and structuring of action execution, with a particular focus on the clustering of behavior into action-reward bouts. We found little effect of enhancing SNc excitability on the proportion of rewards that occurred in action-reward bouts during exploitative states, when animals already perform a high proportion of rewards in action-reward bouts (**Fig. 4f**, bottom row, and **Supp. Fig. 10**). Similarly, we found little difference in this measure when enhancing SNc excitability during exploratory states, before animals had learned that particular action sequences would lead to reward. However, on Day H, when animals could perform a wider range of actions to get the same high rate of reward, we found that enhancing SNc activity led to a significantly higher proportion of the total rewards occurring in action-reward bouts (**Fig. 4f**, bottom row, and **Supp. Fig. 10**, effect was not present in controls expressing mCherry in SNc). The mean number of action-reward pairs per action-reward bout also increased significantly following CNO (**Supp. Fig. 10**). The emergence of this effect only on Day H, when multiple actions can lead to reward, suggested that dopaminergic manipulations might be altering the motivation of action execution in a general manner, rather than targeted towards specific actions alone. We therefore defined “engagement bouts” (see Methods) to quantify the clustering of all task-related actions in time regardless of if they resulted in reward, in contrast to action-reward bouts that only capture the clustering of rewarded actions. We found an increase in the duration of engagement bouts following administration of CNO on Day H (**Fig. 4f**, bottom row, and **Supp. Fig. 10**), suggesting that increasing SNc excitability enhanced the motivation to perform actions broadly in a non-action-specific manner (with high entropy). This restructuring of action-reward bouts and engagement bouts following CNO injections in animals expressing hM3Dq in SNc was not seen in animals expressing hM3Dq in LC (**Supp. Fig. 10**), suggesting that the motivation for frequent action execution is mediated by the dopaminergic, but not noradrenergic, system. Enhanced noradrenergic activity therefore primarily impacts levels of response entropy, driving transitions into exploratory behavioral states and increasing the diversity of actions to be executed, while enhanced dopaminergic activity primarily impacts the motivation to execute actions, resulting in the restructuring of task performance into exploitative bouts of action.

## Discussion

In conclusion, we developed a novel behavioral framework in rodents to capture exploitative and exploratory states of action selection, and we found sustained changes in baseline activity in both dopaminergic and noradrenergic populations across these states. These sustained baseline changes in SNc were due to an increased duration of SNc response transients (plateaus), as well as hysteretic network dynamics, that together resulted in accumulations of activity specifically during exploitative bouts, when animals repeatedly and vigorously performed well-learned actions to achieve the desired outcome. Conversely, the sustained inhibition in LC was caused by a lengthening of negative LC response transients (depressions) during exploitative states, resulting in progressively lower activity levels over the course of exploitative bouts. Importantly, the response dynamics in both regions were strongly influenced not only by the occurrence of individual behavioral events or the macroscopic behavioral state (exploitative or exploratory), but also by the intermediate-scale action-reward bouts nested within these macroscopic states and composed of individual behavioral events. These altered response dynamics in SNc could not be explained by classical RL parameters and instead appeared to correspond to neuronal subpopulations coding for movement vigor, while altered responses in LC corresponded to subpopulations that do not code for movement vigor. Chemogenetic increase of excitability in dopaminergic and noradrenergic populations produced dissociable effects on the motivation and selection of actions, with the dopaminergic system primarily mediating the general motivation to perform actions and the noradrenergic system primarily impacting the diversity of specific actions to be performed. These results uncover two different aspects relating neural mechanisms to behavioral states. The first is that behavioral states are mediated by neural states that can last dozens of seconds, and emerge as a result of subtle changes in the response properties of single neurons in a network, which accumulate over intermediate time scales. The second is that dopaminergic and noradrenergic systems subserve dissociable aspects of exploitative behavioral states.

The response plateaus and depressions observed here with calcium imaging have a number of possible biological underpinnings. Most simply, these altered dynamics could reflect a change in the excitability or gain of individual neurons, which would be consistent with the observed effects following chemogenetic manipulations. This could, as one example, involve a change in the probability of neuronal up− or down-states across the population^21^. Alternatively, these plateaus and depressions could reflect changing lateral interactions within the network, which is also consistent with the observed hysteretic network effects^22^. More granular investigations with physiological methods are necessary to disentangle these possibilities.

The design of our task and experiments allowed us to investigate behavioral and neural states that occur on an intermediate timescale between the scale of synaptic signaling (milliseconds) and the scale of long-term potentiation (hours to days), a timescale that is much more similar to that experienced during ongoing behavior and perhaps more relevant to slower- and longer-acting neuromodulatory systems and the distributed dynamics on which they act. Importantly, there are a number of ways that activity modulations on these intermediate timescales could differentially impact downstream circuits for action selection in the dorsal striatum (DS), in the case of DA, and the anterior cingulate cortex (ACC), in the case of NE. Both systems are known to contain a range of receptor subtypes with distinct postsynaptic effects and behavioral correlates, and these downstream receptors have been hypothesized to respond preferentially to particular temporal dynamics of neuromodulator release^23–28^. Further work is necessary to characterize the changes in target structures produced by neuromodulatory systems during these intermediate-scale behavioral states.

Together, our results suggest that both dopaminergic and noradrenergic signaling modulate the likelihood of transitioning into an exploitative state: a state of inspired engagement in performing a well-known action that might be colloquially referred to as “the hot hand effect” or “being in the zone”. Working in conjunction across multiple temporal scales, these systems structure our execution and selection of behaviors to either maintain a series of successes and capitalize on our learned skills or, conversely, to explore alternative actions and discover novel, creative behavioral responses to an endlessly complex and nuanced environment.

## ACKNOWLEDGMENTS

This material is based on work supported by the National Institutes of Health grant (1K99MH118412-01) to A.C.K., the NARSAD Young Investigator grant (PG010634) to A.C.K., the Simons Collaboration on the Global Brain grant (348880) to A.C.K., the Bial Foundation grant (413/14) to A.C.K., the European Research Council Consolidator grant (COG 617142) to R.M.C., and the U19 Brain Initiative grant (5U19NS104649) to R.M.C. We thank Dr. Vivek Athalye and Dr. James Murray for insightful discussion.

## AUTHOR CONTRIBUTIONS

A.C.K and R.M.C. conceived and designed experiments and analyses. A.C.K performed data collection and analysis. A.C.K. and R.M.C. prepared the manuscript.

## MATERIALS AND METHODS

### Animals

All experiments were performed in compliance with the regulations of the Institutional Animal Care and Use Committee (IACUC) at Columbia University. A total of forty-eight mice (13 female, 35 male) of roughly 3 months of age were used for the experiments. Transgenic mice expressed Cre recombinase under the control of the tyrosine hydroxylase promoter (Tg(Th-cre)FI12Gsat/Mmucd) for targeting of dopaminergic and noradrenergic cells, or Cre recombinase under the control of the dopamine transporter promoter (B6.SJL-Slc6a3tm1.1(cre)Bkmn/J) for targeting of dopaminergic cells.

### Virus Injections

Surgeries were performed under sterile conditions using isoflurane anesthesia (1-3%). Stereotactic coordinates relative to bregma were used to target the SNc (anteroposterior −3.16 mm, mediolateral ±1.4 mm, dorsoventral −4.2 mm) and stereotactic coordinates relative to lambda were used to target the LC (anteroposterior −0.8 mm, mediolateral ±0.8 mm, dorsoventral −3.2 mm). For imaging experiments, animals were injected unilaterally with 500 μL of AAV5.CAG.Flex.GCaMP6f.WPRE.SV40 (University of Pennsylvania Vector Core) into the right SNc or LC. For chemogenetic experiments, experimental animals were injected bilaterally with 500 μL of AAV5-hSyn-DIO-hM3D(Gq)-mCherry (Addgene plasmid #44361), while control animals were injected bilaterally with 500 μL of AAV5-hSyn-DIO-mCherry (Addgene plasmid #50459). All injections were performed using a Nanoject II Injector (Drummond Scientific, Broomall, PA, USA) at a rate of 4.6 nL every 5 seconds. Injection pipettes were left in place for 10 minutes post-injection to allow for virus absorption, and incisions were closed with Vetbond tissue adhesive (3M, Maplewood, MN, USA) for chemogenetic experiments in which no lens was implanted. Animals were given a minimum of 5 days to recover from surgery before behavioral training.

### Chronic Lens Implantation

For imaging experiments, virus injections were followed by implantation of a gradient index (GRIN) lens (Inscopix, Inc., Palo Alto, CA, USA) into the SNc or LC. Overlying tissue was first removed by insertion of a 30-gauge blunt needle to the target site, with care taken to minimize damage. GRIN lenses were then implanted unilaterally and secured to the skull using dental acrylic (Lang Dental, Wheeling, IL, USA). 2-3 weeks were allowed for viral expression before attachment of microendoscope baseplates (Inscopix, Inc.) to the dental acrylic at the correct focal plane for imaging.

### Chemogenetics

For chemogenetic experiments, mice were briefly anesthetized with 1-3% isoflurane and injected intraperitoneally with 5 mg/kg clozapine-N-oxide (CNO) before behavioral sessions. Mice were given 15 minutes following CNO injection to allow for the CNO to take effect and for anesthetic effects to subside.

### Behavioral Task

Animals were trained in custom-made operant boxes (5 in x 6 in) controlled by a python-based framework (PyControl, https://pycontrol.readthedocs.io) that supplies all cues and rewards, as well as recording all behavioral timestamps. Behavior was also monitored with overhead cameras (Flea3, Point Grey Research, Richmond, Canada) recording at 30 frames per second. Operant boxes were placed inside sound attenuating chambers during training. Timestamps from the behavioral task were synchronized with calcium imaging data using TTL pulses sent from the behavioral chambers to the Inscopix data acquisition system.

Operant chambers contained three equidistant nose poke ports surrounding a central reward port. Mice had to discover a rewarded sequence of three pokes in a specific order with no intervening pokes. Importantly, the task contains no trial structure and few cues, ensuring that mice actively explore the environment to discover what is rewarding. When a correct sequence was performed, water rewards of 5-15 μL were supplied through the opening of a solenoid.

Mice were initially pre-trained in a setting in which any possible three-poke sequence that includes all three nose poke ports was rewarded. Following roughly one week of pre-training, mice were exposed to the full task, in which only one target sequence was rewarded. Once mice achieved proficiency on a particular target sequence, the rewarded sequence was changed. For experiments on Day H, all three-poke sequences were rewarded with 80% probability. For experiments on Day L, all three-poke sequences were rewarded with 20% probability. In all cases, rewards could not be cached and had to be consumed prior to earning further rewards.

During calcium imaging experiments, fluorescence images were acquired at a frame rate of 10 hz.

### Data Analysis

Analyses were performed in Matlab (Mathworks, Natick, MA) with custom-written routines. Behavioral data were sampled in 1 ms bins. For sliding window analyses of behavioral data, a window size of 100 trials with a step size of 5 trials was used. For each behavioral session, histograms were created for the empirical probability of performing each sequence (“Sequence Entropy”), as well as for the empirical probability of transitions between nose pokes (“Transition Entropy”), and entropy was calculated as:

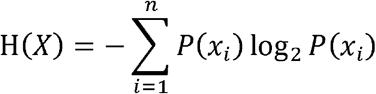

Corrections for finite sample sizes^13,42^ were tested by sampling from a known distribution with a structure similar to that seen in our behavioral data, and these corrections were found to be less accurate than the above formula in measuring the entropy of the parent distribution. These corrections were therefore not used in subsequent analyses.

A basic reinforcement learning (RL) model was also applied to the behavioral data. Expected action values, Q, were updated on each trial according to:

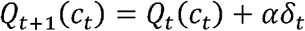

where *c_t_* is the choice on trial t, *δ_t_* is the reward prediction error on trial t, and *α* is the learning rate of the model. Expected action values were related to choices by the following equation:

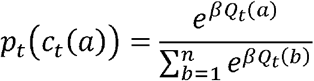

where *β* is the inverse temperature parameter. Finally, expected state values, V, were estimated as the sum of all current action values in that state weighted by their probability of occurrence:

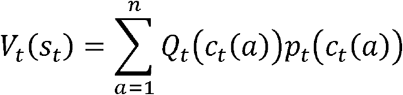

The learning rate and inverse temperature were fit using maximum likelihood estimation.

Behavioral data were also separated into “action-reward bouts” and “engagement bouts”. “Action-reward bouts” were defined as groups of multiple action-reward events (where animals performed the target action sequence and were rewarded for it) that were separated from each other by less than 10 seconds and separated from other action-reward events by more than 20 seconds. “Engagement bouts”, on the other hand, were defined as groups of multiple individual actions (single task pokes, rather than target three-poke sequences) that were separated from each other by less than 2 seconds and separated from other actions by more than 5 seconds, irrespective of whether or not the actions resulted in reward.

Calcium imaging data were first preprocessed using Mosaic (Inscopix, Inc.) to apply 4x spatial downsampling and motion correction. Constrained non-negative matrix factorization (CNMF-E)^29–30^ was then applied for denoising and demixing of the data. The footprints and activity profiles of all putative neurons were inspected manually before inclusion in the dataset. 121 cells in SNc, 61 cells in LC, and 9 cells in VTA were included in the primary analyses. Unless otherwise noted, significance was assessed at the p < 0.05 level using a two-tailed T-test, with bonferroni corrections applied when multiple comparisons were performed.

For the extraction of quickly-varying components of fluorescence signals, dF/F was calculated on these traces with a relatively short sliding window of 5 seconds. For the extraction of slowly-varying components of the fluorescence signals, traces were smoothed with a moving average of 60 seconds.

Occurrences of reward bouts were predicted from neural data using a wiener filter. Five lags were used occurring every 500 milliseconds starting 2 seconds before bout start and ending at bout start. Prediction was done for each cell individually to assess their contribution. Prediction performance was assessed using the area-under-the-curve (AUC) from a receiver operating characteristic (ROC) curve. Results were also compared to results obtained when the behavioral category labels were shuffled.

To separate vigor-correlated and non-vigor-correlated neuronal populations, a median split was performed on the data based on the correlation coefficient with the negative of the inter-poke interval. The neurons most correlated with fast, vigorous poking were considered vigor-correlated neurons and all others were considered non-vigor-correlated.

For convolution models, pure exponentials of varying lengths were used to model the SNc impulse response function. For the LC impulse response function, the average response from all LC cells to unexpected rewards was smoothed by a 1 second moving average. To model hysteretic network dynamics, multiple convolution traces were created for each animal with the addition of a random temporal jitter in the response to reward for each trace that preferentially shifts responses positively in time by a random fraction of a maximum of 5 seconds.

To quantify response transient durations, positive and negative threshold crossings (3SD) were located. A 5 second window before threshold crossing was defined as baseline activity, and a 5 second window after threshold crossing was then advanced until the average activity in this window matched the average activity in the baseline window. The number of timepoints by which the second window had to be advanced to equal the baseline window was defined as the response transient duration. Individual transients were also fit with exponential functions to determine decay and offset parameters.

## SUPPLEMENTARY FIGURES

**Supplementary Figure 1.**
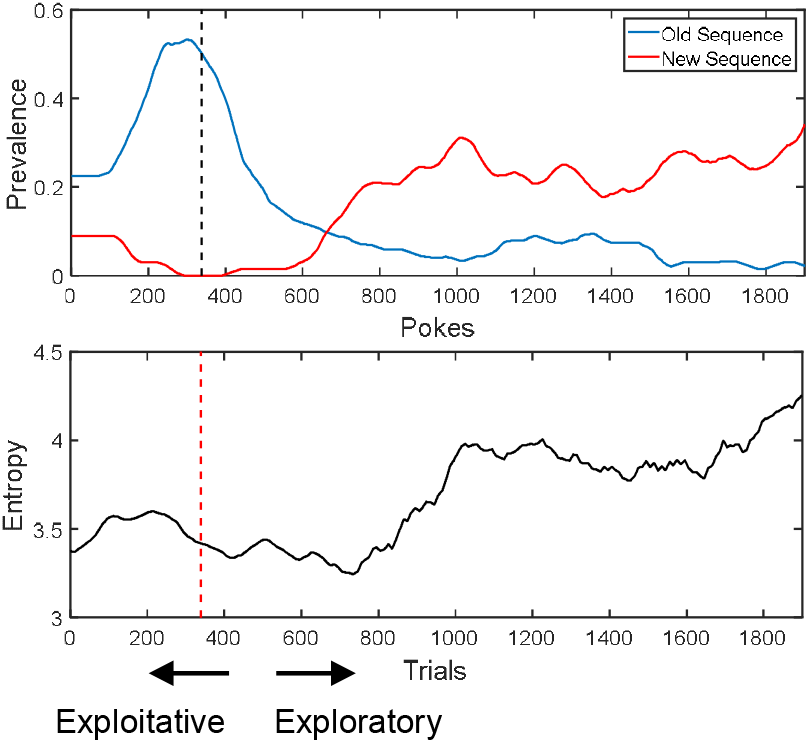
Sequence prevalence and choice entropy surrounding a change in the rewarded sequence. Animals begin reversal sessions exploiting a well-known sequence and the rewarded sequence is changed during the session. Top: Prevalence of the sequence that was rewarded before (blue) or after (red) the change in rewarded sequence (dotted line). Bottom: The entropy of selected actions rises after the change in rewarded sequence (dotted line) and falls again as animals discover the new rewarded sequence.

**Supplementary Figure 2.**
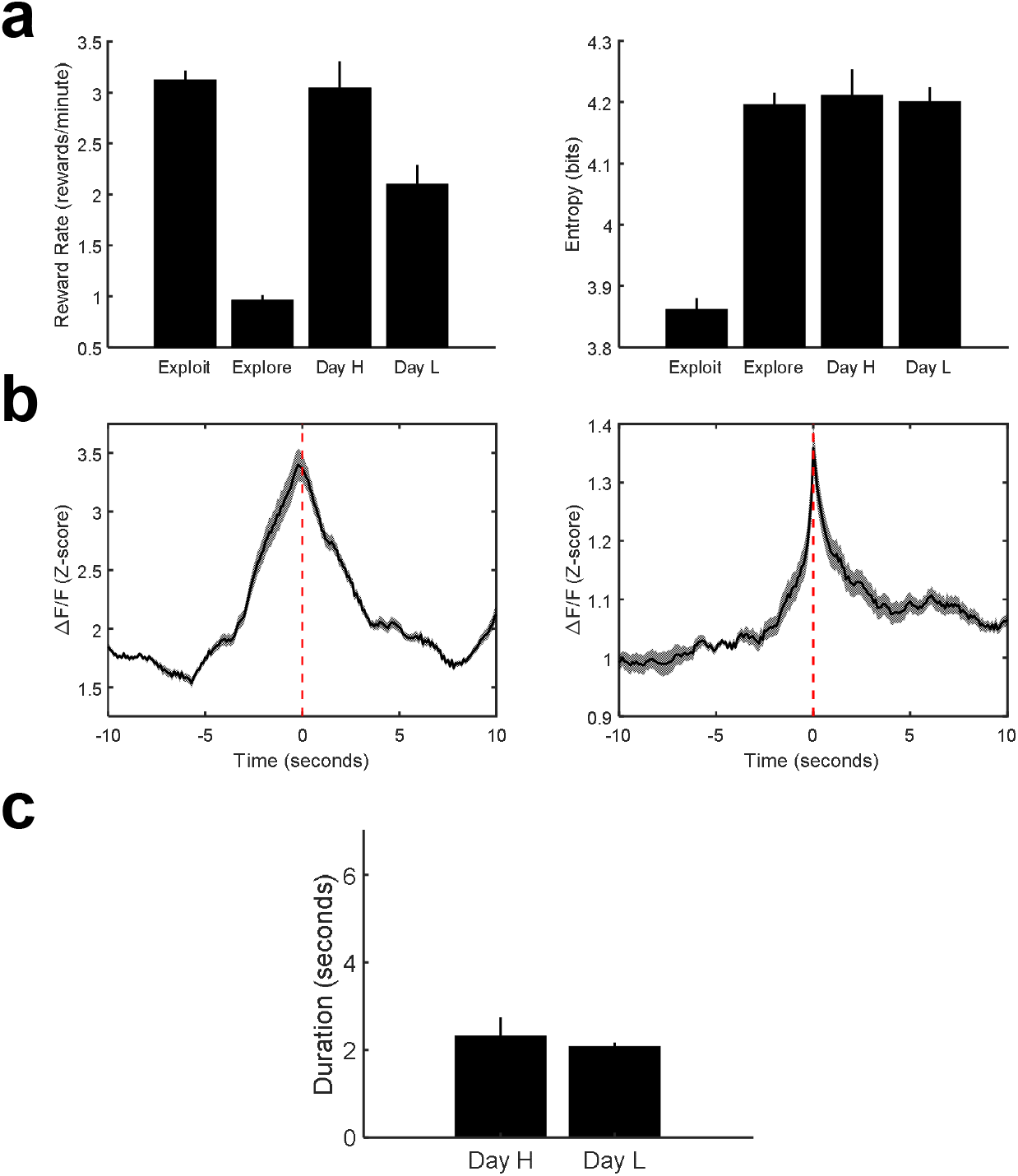
Behavioral and neural metrics on Day H and Day L. **a.** Reward rate (left) and response entropy (right) during exploitation, exploration, on Day H, and on Day L. **b.** Cross-correlation histograms of activity in SNc time-locked to large fluorescence bursts in other cells in the network on Day H (left) and Day L (right). **c.** Duration of positive transients on Day H and Day L.

**Supplementary Figure 3.**
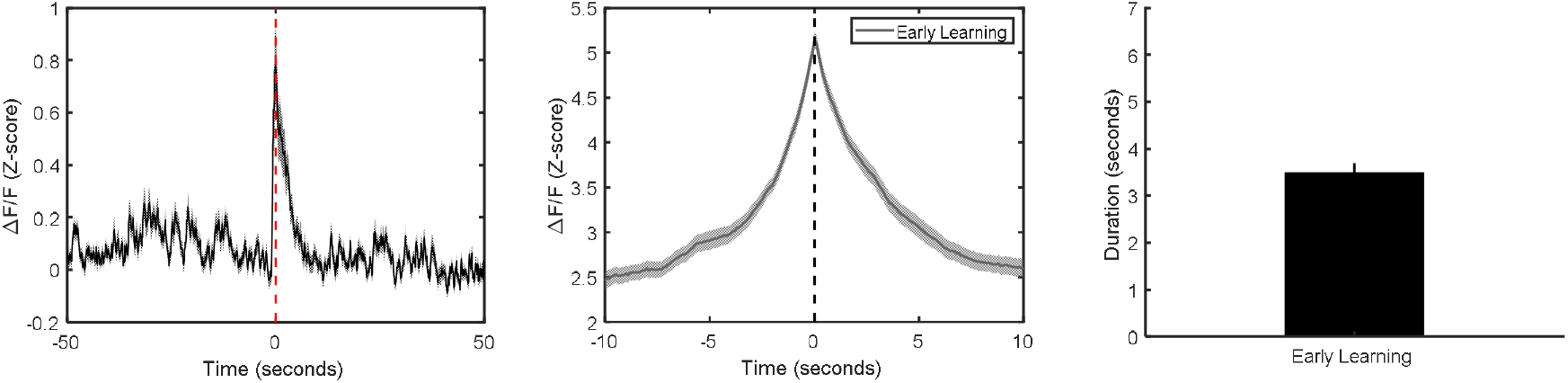
SNc effects are not present early in training. PSTH of SNc activity time-locked to exploitative rewards (Left), cross-correlation histogram (center), and duration of positive transients (right) during early training.

**Supplementary Figure 4.**
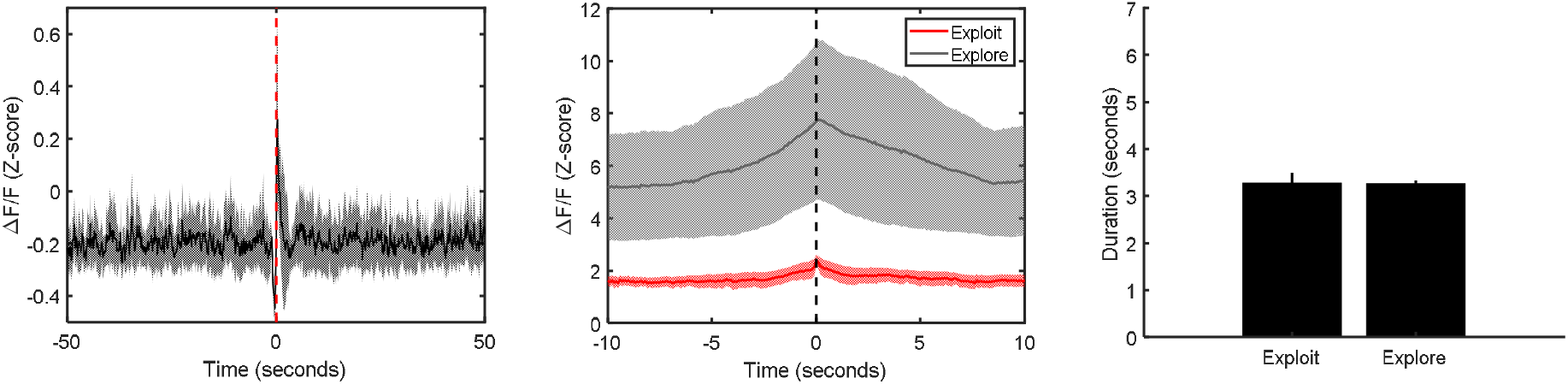
Effects observed in SNc are not present in VTA. PSTH of VTA activity time-locked to rewards (Left), cross-correlation histogram (center), and duration of positive transients (right) during exploitative states.

**Supplementary Figure 5.**
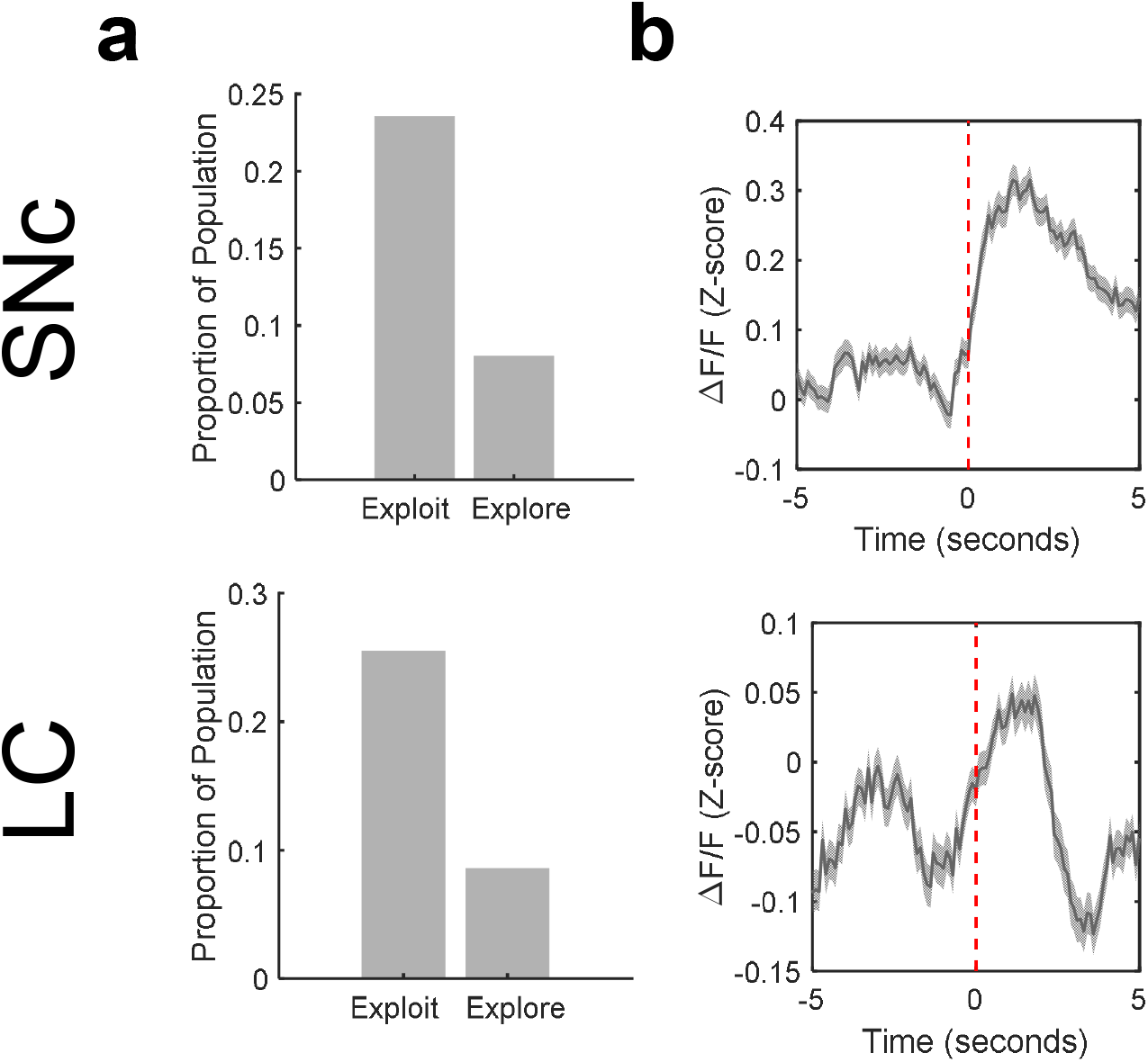
A subpopulation of the network exhibits sustained activity and codes for movement vigor. **a.** Mean proportion of the network exhibiting sustained activity in SNc (top) and LC (bottom) during exploitative or exploratory behavioral states. **b.** Mean response of the network time-locked to initiation of the rewarded action in SNc (top) and LC (bottom). The neuronal impulse response functions in both regions align closely with this behavioral event, whereas they align less well to reward achievement (Fig. 1k). Error bars denote s.e.m.

**Supplementary Figure 6.**
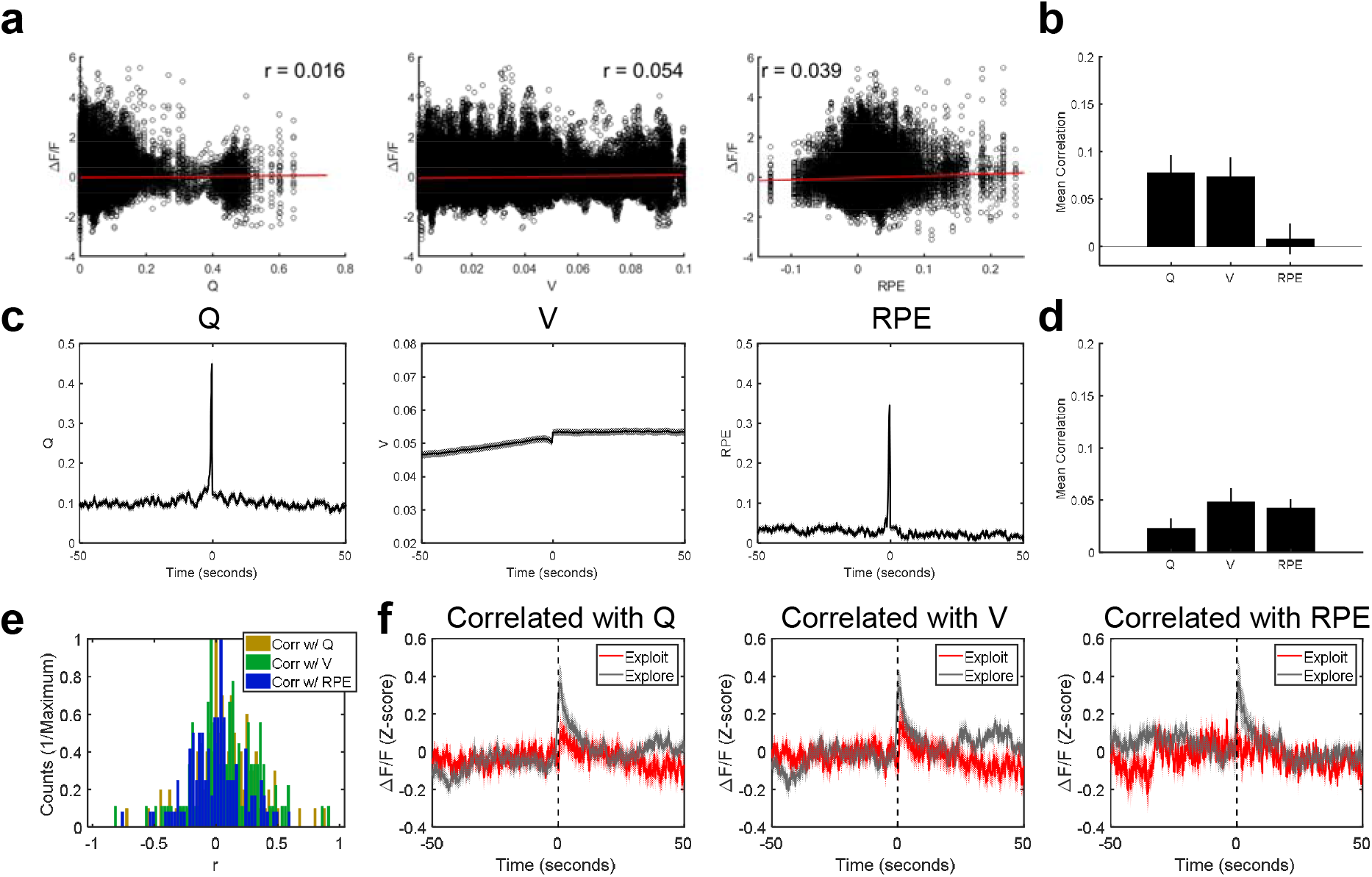
SNc effects are not explained by basic RL parameters. **a.** Scatter plots showing mean neuronal activity within 2 seconds of an action versus that trial’s current estimate of action value (left), state value (center), and reward prediction error (right). **b.** Mean correlations of peri-event neuronal activity in individual cells with action-by-action estimates of action value, state value, and reward prediction error. **c.** PSTHs time-locked to reward using action-by-action estimates of current action value (Q, left), state value (V, center), or reward prediction error (RPE, right). **d.** Mean correlation of full session time courses of action value, state value, and reward prediction error with smoothed SNc neuronal activity. **e.** Individual neurons exhibit a range of correlations with RL estimates of action value (black), state value (red), or RPE (blue). **f.** Neurons with positive activity correlations (r>0.1 and p<0.05) with either action value (left), state value (middle), or RPE (right) do not exhibit increased levels of sustained activity during Exploit relative to Explore-New conditions. These subsets of the population exhibit decreases in phasic responses during exploitation (when rewards are expected) relative to exploration, in agreement with classic reports of RPE coding in dopaminergic neurons, but notably distinct from other recorded neurons in the population. Error bars denote s.e.m.

**Supplementary Figure 7.**
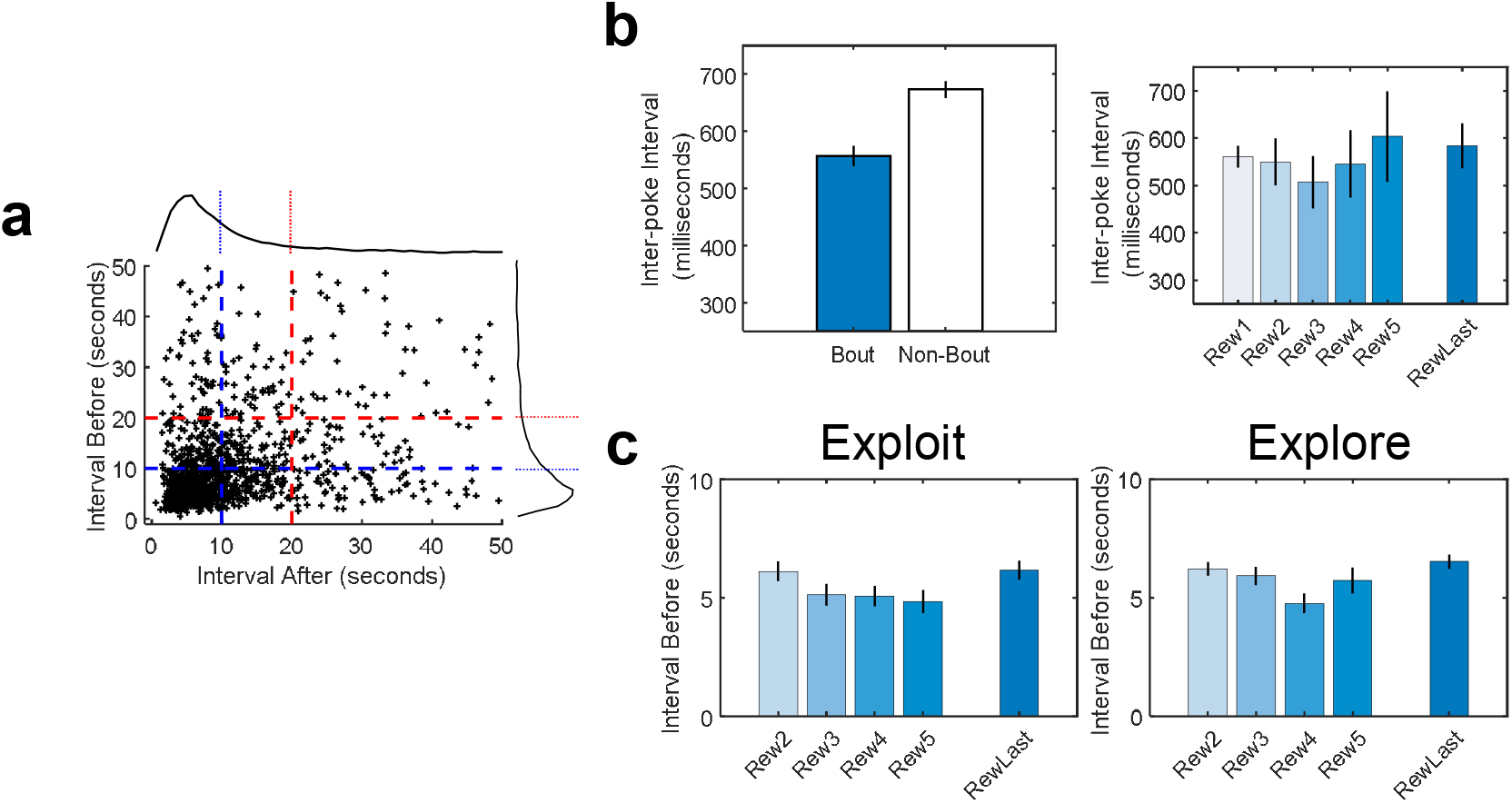
Characteristics of Action-Reward Bouts. **a.** Scatter plot showing the interval before versus the interval after each reward. Many bouts of action-reward pairs occur within 10 seconds of each other, while far fewer occur in intervals greater than 20 seconds. **b.** The inter-poke interval is significantly faster for pokes leading to reward during exploitative action-reward bouts than it is for pokes leading to reward outside of action-reward bouts (left), but does not vary based on position within an action-reward bout (right). **c.** The inter-poke interval within a bout is not significantly different during exploitative (left) and exploratory (right) states or based on position within the bout.

**Supplementary Figure 8.**
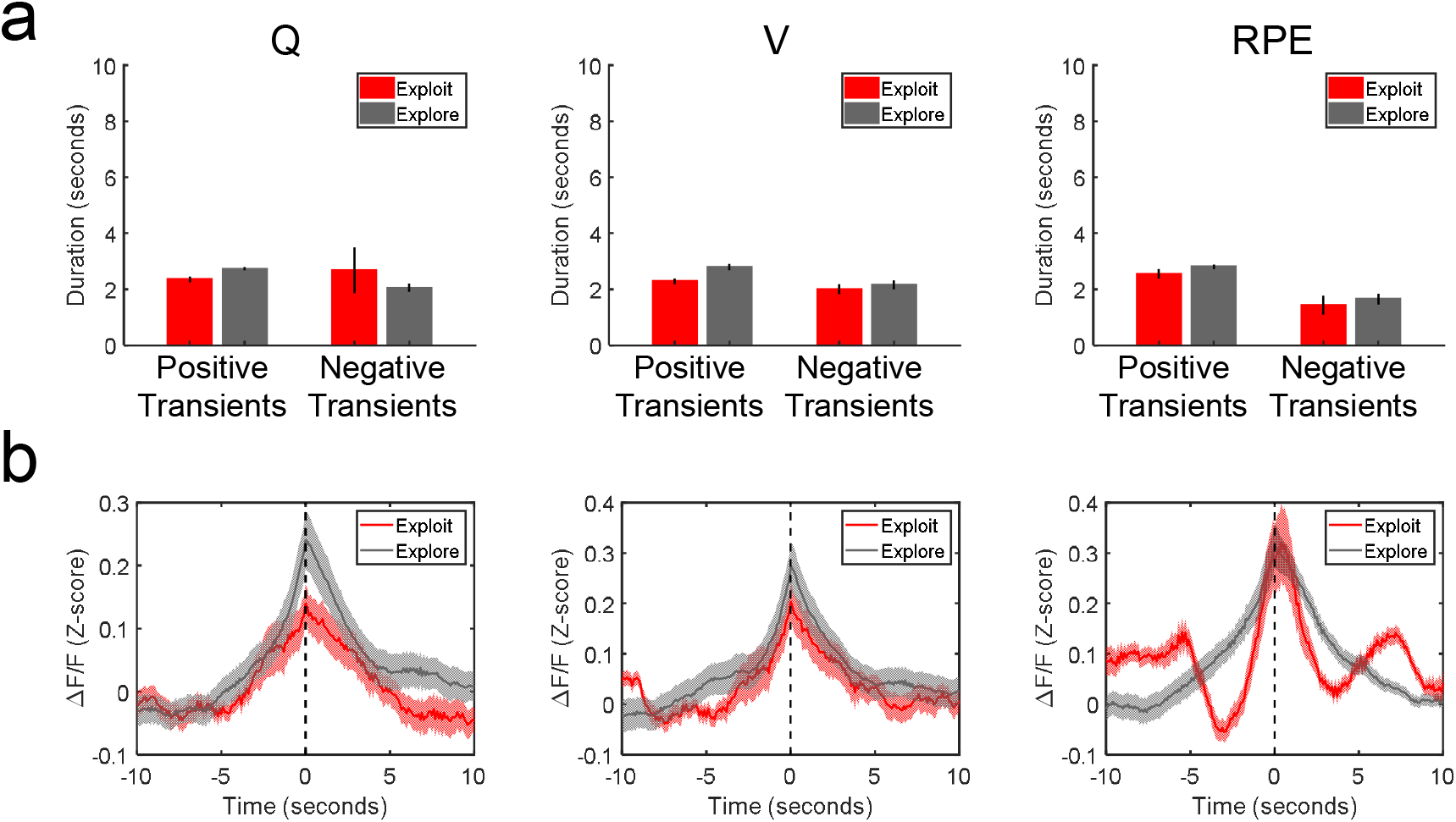
SNc cells that are correlated with RL parameters do not exhibit plateaus or hysteresis. **a.** Length of positive and negative transients for cells correlated with action value (left), state value (middle), or RPE (right). While the observed sustained effects were found to be due to lengthened response plateaus in SNc during exploitative states, cells exhibiting correlations with RL parameters instead show shorter positive transients during exploitative (red) relative to exploratory (gray) states. Transient durations are generally low for these sub-populations. **b.** Cross-correlation histograms for cells correlated with action value (left), state value (middle), or RPE (right). This population does not exhibit the asymmetrical hysteretic network effects observed in the general SNc population during exploitative relative to exploratory states. Error bars denote s.e.m.

**Supplementary Figure 9.**
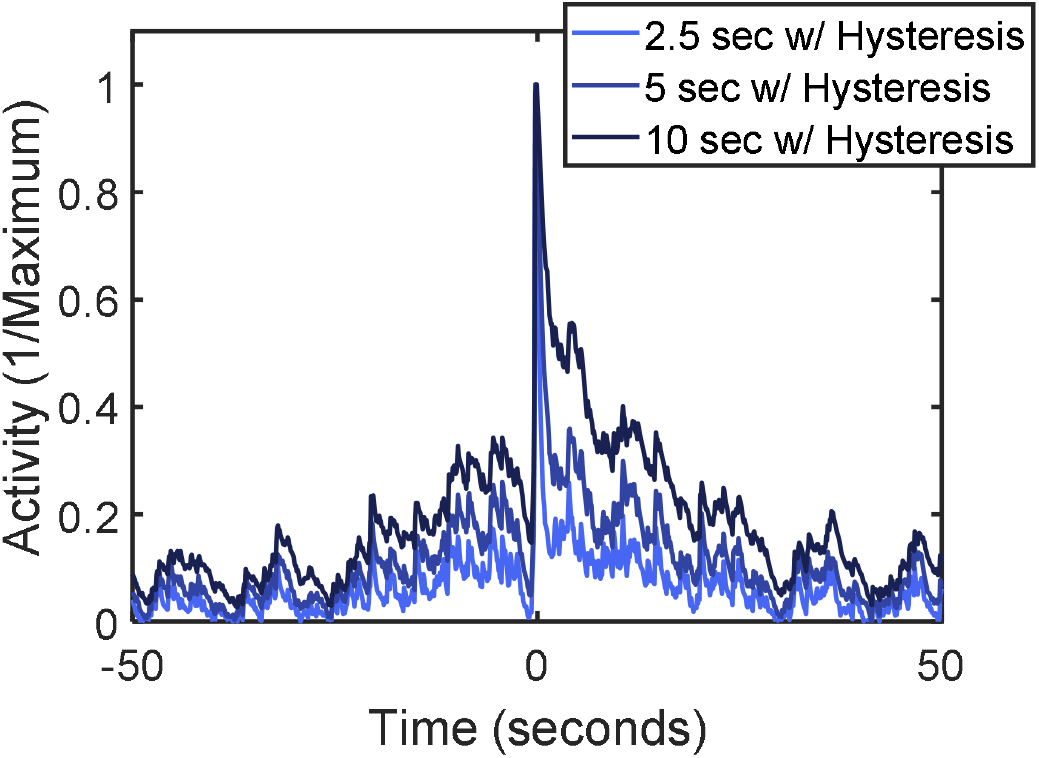
Reward convolution model requires both lengthened IRF and hysteresis to recapitulate effects. Including hysteresis in the reward convolution model alters the response shape for all IRFs, but is not sufficient to recapitulate the effects observed in our data without also including a lengthened IRF. Hysteresis with short IRFs does not replicate the data.

**Supplementary Figure 10.**
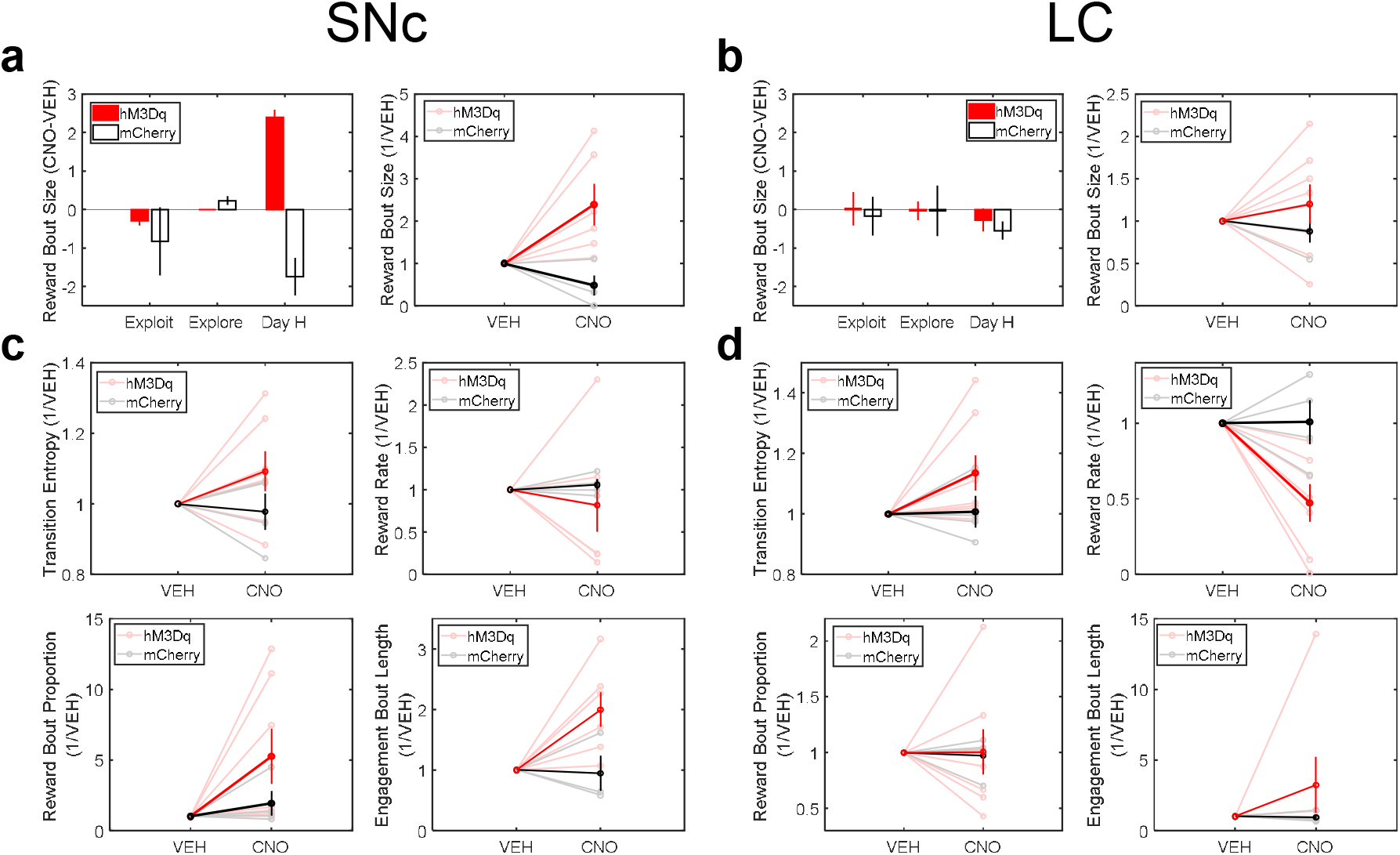
Chemogenetic effects are specific to dopaminergic and noradrenergic systems. **a-b.** The mean number of action-reward pairs per action-reward bout in animals expressing hM3Dq (red) or mCherry (black) in SNc (a) or LC (b). Left: Bar graphs depicting means for all conditions. Right: Line plots depicting the change in individual animals following VEH or CNO injections, normalized to values seen following VEH injections. **c-d.** The transition entropy (top left), reward rate (top right), proportion of action-reward pairs that occurred in bouts (bottom left), and mean duration of engagement bouts (bottom right), shown for individual animals expressing hM3Dq (red) or mCherry (black) in SNc (c) or LC (d), normalized to values seen following VEH injections. Error bars denote s.e.m. Light traces represent individual animals, dark traces represent population means.

## REFERENCES

1. Costa, R.M. (2007). Plastic corticostriatal circuits for action learning: what’s dopamine got to do with it? Ann NY Acad Sci, 1104:172–91.

2. Aston-Jones, G. & Cohen, J.D. (2005). An integrative theory of locus coeruleus-norepinephrine function: adaptive gain and optimal performance. Annu Rev Neurosci, 28:403–450.

3. Usher, M., Cohen, J.D., Servan-Schreiber, D., Rajkowski, J., & Aston-Jones, G. (1999). The role of locus coeruleus in the regulation of cognitive performance. Science, 283:549–554.

4. Izquierdo, A., Wiedholz, L.M., Millstein, R.A., Yang, R.J., Bussey, T.J., Saksida, L.M., & Holmes, A (2006). Genetic and dopaminergic modulation of reversal learning in a touchscreen-based operant procedure for mice. Brain Behav Res, 171:181–88.

5. Klanker, M., Feenstra, M., & Denys, D. (2013). Dopaminergic control of cognitive flexibility in humans and animals. Front Neurosci, 7:201.

6. Humphries, M.D., Khamassi, M., & Gurney, K. (2012). Dopaminergic control of the exploration-exploitation trade-off via the basal ganglia. Front Neurosci, doi: 10.3389/fnins.2012.00009.

7. Tervo, D.G.R., Proskurin, M., Manakov, M., Kabra, M., Vollmer, A., Branson, K., & Karpova, A.Y. (2014). Behavioral variability through stochastic choice and its gating by anterior cingulate cortex. Cell, 159(1):21–32.

8. Karlsson, M.P., Tervo, D.G., & Karpova, A.Y. (2012). Network resets in medial prefrontal cortex mark the onset of behavioral uncertainty. Science, 338(6103):135–9.

9. Cohen, J.D., McClure, S.M., & Yu, A.J. (2007). Should I stay of should I go? How the human brain manages the trade-off between exploitation and exploration. Philos Trans R Soc Lond B Biol Soc, 362:933–942.

10. Dayan, P. & Daw, N.D. (2008). Decision theory, reinforcement learning, and the brain. Cogn Affect Behav Neurosci, 8:429–53.

11. Daw, N.D., Niv, Y., & Dayan, P. (2005). Uncertainty-based competition between prefrontal and dorsolateral striatal systems for behavioral control. Nat Neurosci, 8:1704–11.

12. Ebitz, R.B., Albarran, E., & Moore, T. (2018). Exploration disrupts choice-predictive signals and alters dynamics in prefrontal cortex. Neuron, 97(2):450–61.

13. Speekenbrink, M. & Konstantinidis, E. (2015). Uncertainty and exploration in a restless bandit problem. Top Cogn Sci, 7:351–67.

14. Neiman, T. & Loewenstein, Y. (2011). Reinforcement learning in professional basketball players. Nat Commun, 2:569.

15. Blanchard, T.C., Wilke, A., & Hayden, B.Y. (2014). Hot-hand bias in rhesus monkeys. J Exp Psychol Anim Learn Cogn, 40(3):280–6.

16. Hamid, A.A., Pettibone, J.R., Mabrouk, O.S., Hetrick, V.L., Schmidt, R., Vander Weele, C.M., Kennedy, R.T., Aragona, B.J., & Berke, J.D. (2016). Mesolimbic dopamine signals the value of work. Nat Neurosci, 19(1):117–126.

17. da Silva, J.A., Tecuapetla, F., Paixao, V., & Costa, R.M. (2018). Dopamine neuron activity before action initiation gates and invigorates future movements. Nature, 554(7691):244–248.

18. Schultz, W., Dayan, P., & Montague, P.R. (1997). A neural substrate of prediction and reward. Science, 275(5306):1593–9.

19. Armbruster, B.N., Li, X., Pausch, M.H., Herlitze, S., & Roth, B.L. (2007). Evolving the lock to fit the key to create a family of G protein-coupled receptors potently activated by an inert ligand. Proc Natl Acad Sci, 104(12):5163–8.

20. Urban, D.J. & Roth, B.L. (2015). DREADDs (designer receptors exclusively activated by designer drugs): chemogenetic tools with therapeutic utility. Anny Rev Pharmacol Toxicol, 55:339–417.

21. UP-DOWN cortical dynamics reflect state transitions in a bistable network. Elife, 6: doi: 10.7554/eLife.22425.

22. Sahasranamam, A., Vlachos, I., Aertsen, A., & Kumar, A. (2016). Dynamical state of the network determines the efficacy of single neuron properties in shaping the network activity. Sci Rep, 6:26029.

23. Kravitz, A.V., Tye, L.D., & Kreitzer, A.C. (2012). Distinct roles for direct and indirect pathway striatal neurons in reinforcement. Nat Neurosci, 15(6):816–8.

24. Tecuapetla, F., Jin, X., Lima, S.Q., & Costa, R.M. (2016). Complementary contributions of striatal projection pathways to action initiation and execution. Cell, 166(3):703–715.

25. Dreyer, J.K., Herrik, K.F., Berg, R.W., & Hounsgaard, J.D. (2010). Influence of phasic and tonic dopamine release on receptor activation. J Neurosci, 30:14273–83.

26. Berridge, C.W. & Spencer, R.C. (2016). Differential cognitive actions of norepinephrine a2 and a1 receptor signaling in the prefrontal cortex. Brain Res, 1641: 189–96.

27. Lohani, S., Martig, A.K., Deisseroth, K., Witten, I.B., Moghaddam, B. (2019). Dopamine modulation of prefrontal cortex activity is manifold and operates at multiple temporal and spatial scales. Cell Rep, 27(1):99–114.

28. Arnsten, A.F. (2000). Through the looking glass: differential noradrenergic modulation of prefrontal cortical function. Neural Plast, 7(1-2):133–46.

29. Zhou, P., Resendez, S.L., Rodriguez, Romaguera, J., Jimenez, J.C., Neufeld, S.Q., Giovanucci, A., Friedrich, J., Pnevmatikakis, E.A., Stuber, G.D., Hen, R., Kheirbek, M.A., Sabatini, B.L., Kass, R.E., & Paninski, L. (2018). Efficient and accurate extraction of in vivo calcium signals from microendoscopic video data. Elife, doi: 10.7554/eLife.28728.

30. Pnevmatikakis, E.A., Soudry, D., Gao, Y., Machado, T.A., Merel, J., Pfau, D., Reardon, T., Mu, Y., Lacefield, C., Yang, W., Ahrens, M., Bruno, R., Jessell, T.M., Peterka, D.S., Yuste, R., &Paninski, L. (2016). Simultaneous denoising, deconvolution, and demixing of calcium imaging data. Neuron, 89(2):285–99.

